# Fabrication of sharp silicon arrays to wound *Caenorhabditis elegans*

**DOI:** 10.1101/776302

**Authors:** Jérôme Belougne, Igor Ozerov, Céline Caillard, Frédéric Bedu, Jonathan J. Ewbank

## Abstract

Understanding how animals respond to injury and how wounds heal remains a challenge. These questions can be addressed using genetically tractable animals, including the nematode *Caenorhabditis elegans*. Given its small size, the current methods for inflicting wounds in a controlled manner are demanding. To facilitate and accelerate the procedure, we fabricated regular arrays of pyramidal features (“pins”) sharp enough to pierce the tough nematode cuticle. The pyramids were made from monocrystalline silicon wafers that were micro-structured using optical lithography and alkaline wet etching. The fabrication protocol and the geometry of the pins, determined by electron microscopy, are described in detail. Upon wounding, *C. elegans* expresses genes encoding antimicrobial peptides. A comparison of the induction of antimicrobial peptide gene expression using traditional needles and the pin arrays demonstrates the utility of this new method.

## Introduction

As part of the investigation of innate immune defences in *C. elegans*, worms are generally wounded in one of two ways. For precise cell-biological studies, individual worms mounted on glass slides can be injured using a laser coupled with an inverted microscope. As a complementary method, worms can be wounded manually, using standard glass microinjection needles^1^. Both techniques are time-consuming and require meticulousness. More than 20 years ago, a method for genetically transforming nematodes using arrays of silicon micromechanical piercing structures was published^2, 3^. This inspired us to investigate their possible use as a tool to wound *C. elegans* in a less painstaking manner.

Silicon is the most widely used material in the microelectronics industry and in the production of micro-electromechanical systems. It is abundant, inexpensive and mechanically robust^4^. High-quality silicon wafers are widely available commercially and fabrication processes for silicon features have been developed for more than 50 years^5^. Silicon microstructures have been already successfully used for piercing of biological objects^2^ and for the fabrication of microneedles^6^ for gene and drug delivery^7^. These studies have shown that silicon is a chemically and mechanically stable biocompatible material. Despite the apparent simplicity of the method described by Trimmer and colleagues^2^, there appears to be no other study using these arrays to transform any nematode species. On the other hand, such arrays could be well suited for use as tools to wound worms^8^. Unfortunately, neither the arrays nor a protocol for their fabrication are available any more (W. Trimmer, personal communication). In order to fabricate equivalent microstructures we therefore developed our own protocol, based on optical lithography and alkaline wet etching.

## Materials and Methods

### Array production

As initial material, we used n-type monocrystalline silicon wafers doped by phosphorus (5 − 10 Ω*cm*) oriented with [100] as the surface plane (from Sil’tronix, France). The wafers were polished on both sides and covered with a 290 nm–thick thermally grown oxide layer. We used two sizes of wafers with diameters of 1 inch (25.4 ± 0.3*mm*) and 4 inches (100 ± 0.3*mm*).

All the microfabrication processes were carried out in a clean room of air purity class ISO 5. The wafers were cleaned in successive baths of acetone and isopropyl alcohol (IPA) with ultrasonic agitation, then rinsed with deionized water and dried under clean nitrogen flow. In order to improve the adhesion of the photoresist and to remove the traces of organic solvents and humidity, the wafers were exposed to oxygen plasma at a temperature of 150°*C* for 10 minutes in a plasma chamber (Nanoplas, France). The radiofrequency power for plasma excitation was 200W. Then, the wafers were spin-coated with a positive photoresist (Microposit S1813 from MicroChem) and exposed to UV-light using MJB-4 (Suss MicroTec, Germany) or PLA-501 (Canon, Japan) optical lithography tools through a mask of round features.

We studied two types of arrays using masks with different geometries: hexagonal and square arrays. The hexagonal arrays were denser than the square ones, but in both cases we kept the distance between the features equal to 100 *μm* in order to prevent possible proximity effects. During the lithography step, we usually oriented the x-axis of the arrays along the [110] silicon crystalline axis in order to cut the samples in between the features during the silicon cleavage; the orientation should not affect the final shape of individual microfeatures^9^.

The feature diameters were 75, 100 and 300 *μm* and the center-to-center distances were 100, 133 and 400 *μm* respectively in the hexagonal arrays. The mask designs for the 75 and 100 *μm* feature arrays are shown in Fig. 1. For square arrays, we used only 300 *μm* features with 400 and 600 *μm* center-to-center distances (pitch). After UV light exposure the features were developed in a commercially available solution (Microposit MF-319) containing tetramethylammonium hydroxide (TMAH). The thickness of the deposited photoresist was about 1.4 *μm* and it was monitored by a mechanical contact profilometer (Dektak, Bruker).

**Figure 1.**
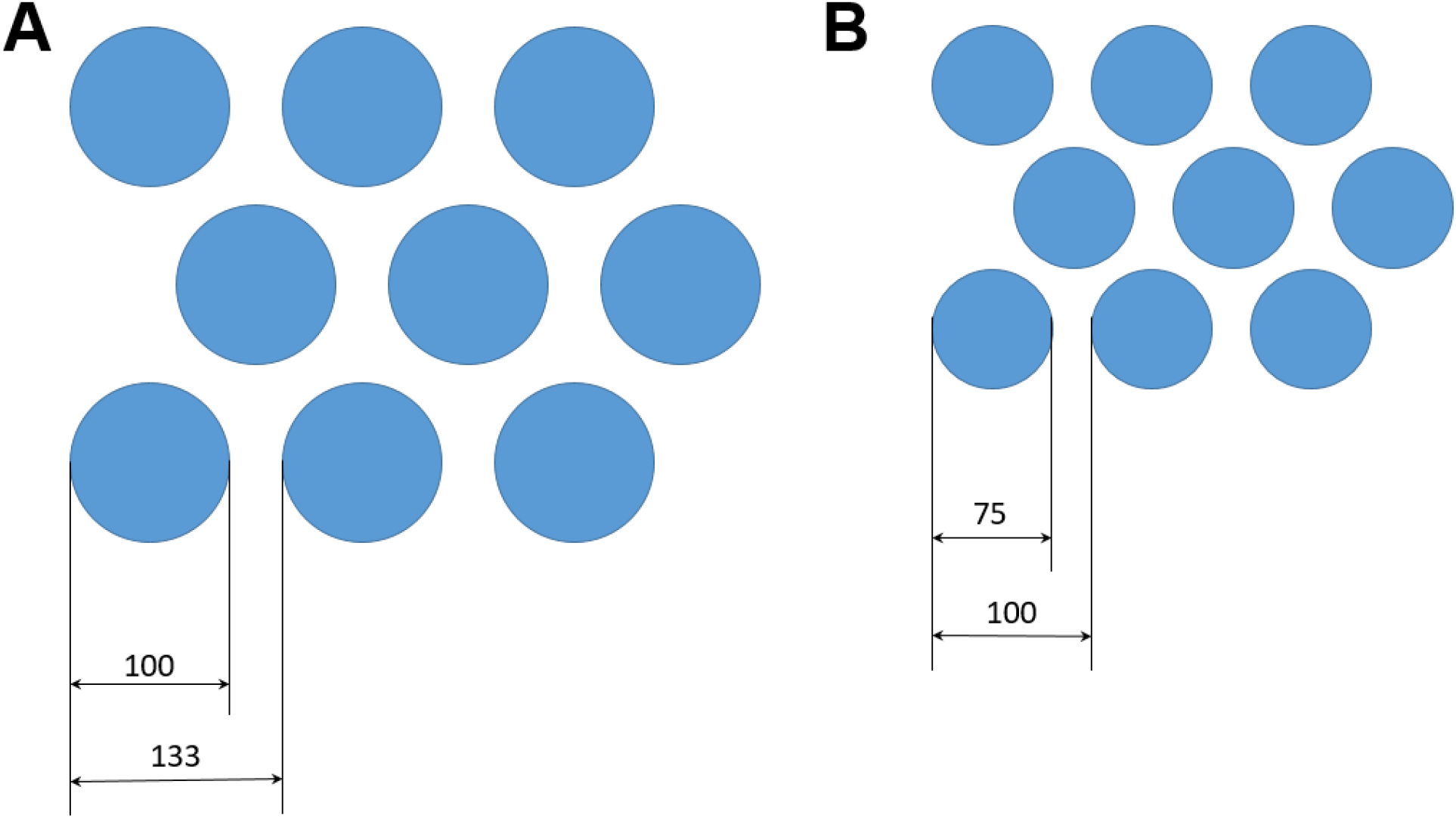
Design of the pattern on the photolithography mask. The round features correspond to the dark (chromium coated) areas on the mask. A) 100 *μ*m hexagonal array; B) 75 *μ*m hexagonal array.

The features were transferred from the resist layer to silicon dioxide by reactive ion etching (MG-200, Plassys, France). The silicon dioxide was completely removed from the topside of the wafers in the areas between the features that were protected by photoresist. We used CHF_3_ reactive plasma because of its good selectivity in etching of SiO_2_, silicon and the photoresist polymer^10^. The remaining photoresist was removed in a solution of 1165 (from Microposit) containing N-methyl-2-pyrrolidone. We confirmed complete removal of the oxide layer in the areas between the round features by contact profilometry. Finally, several wafers were directly processed by alkaline solution while the other wafers were cut in 1 cm × 1 cm square pieces along [110] the crystalline directions of the silicon crystal using a diamond scriber.

Each sample was exposed to potassium hydroxide (KOH) solution in order to etch silicon in the areas not protected by the SiO_2_ hard mask. Wet chemical attack by hydroxide solutions is known to be anisotropic because KOH etches different crystalline planes with significantly different rates^11^. We chose a relatively high KOH concentration of 45% (in deionized water; Fluka) because the resulting surface quality is known to be better for highly concentrated solutions (see page 40 in Fruhauf *et al.*^5^). In several experimental series, we also saturated the solution by addition of isopropanol (IPA) or ethanol.

A borosilicate glass beaker containing the etching solution was placed into a water bath on a regulated hotplate. The etching solution was kept at a temperature of 60 – 75°*C* during the etching in order to control the shape of the obtained features, which was also influenced by the etching time. The temperature was measured by a thermometer immersed in the KOH solution. We developed a series of custom designed magnetic 3-D printed sample holders allowing constant uniform agitation for the samples of different sizes and shapes^12^. The samples were fixed on the magnetic holders fabricated from a temperature resistant photopolymer plastic resin (Formlabs High Temp Resin) and then immersed in the etching solution. Before their first use, the sample holders were rinsed in a hot KOH solution for at least one hour, to remove polymer residues that otherwise interfered with alkaline etching. The samples were agitated at high speed in hot alkaline solution on a standard bench-top magnetic stirrer in order to homogenize the etching and to disperse hydrogen bubbles that appear during the chemical reaction of silicon with hydroxide ions^11^. It was possible to install up to four small samples onto the holders (Figure 2) and to process them in the same etching run. The samples could be then extracted one by one after the desired etching times. The 4 inch wafers were processed one-by-one. In order to keep the most stable conditions we used the same quantity of KOH (650-750 ml) in the beaker and we changed the solution regularly to prevent the alteration of its composition through the introduction of chemical reaction products and by evaporation.

**Figure 2.**
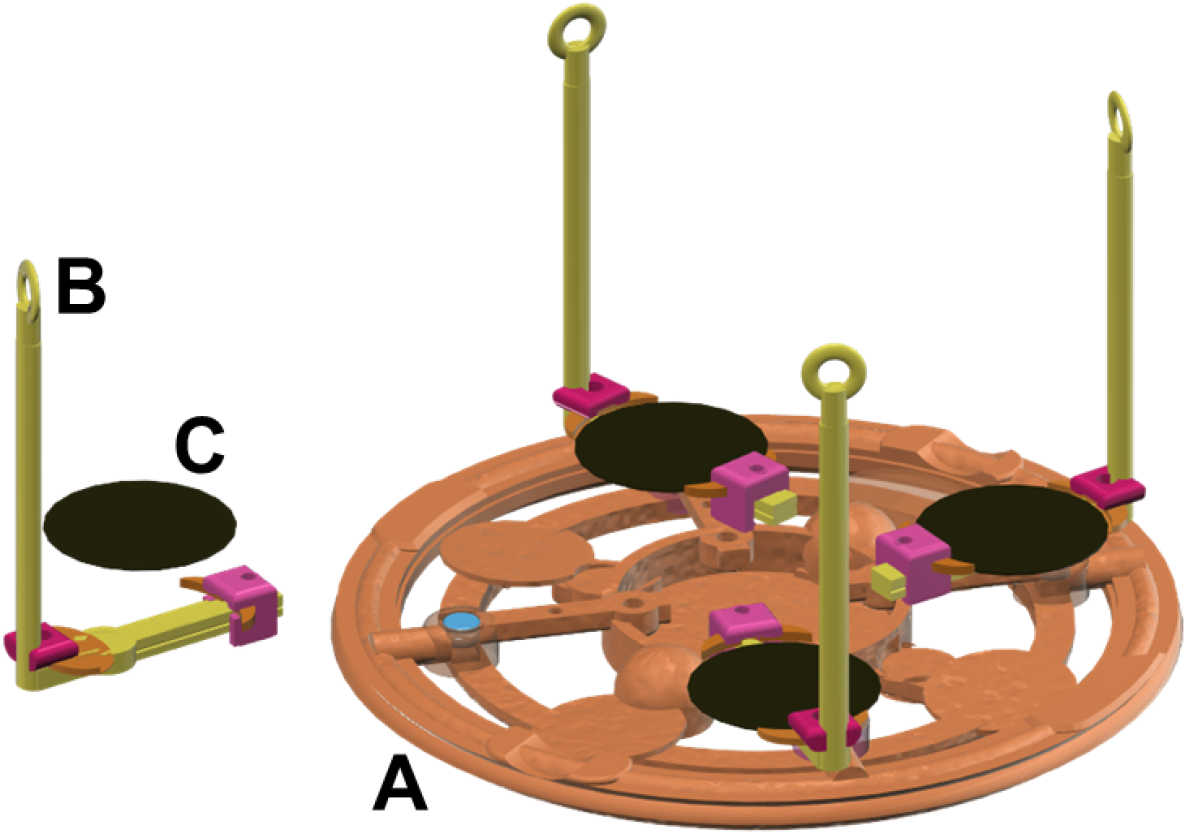
Drawing of a sample holder used for homogeneous KOH etching of multiple one-inch wafers (Autodesk 123 Design software). A) Main support with magnets; B) Arms for easy removal from etching solution; C) One-inch (2.54 cm) diameter wafer to be etched

The typical etching time in KOH solution depended on the feature size. For the 75 *μ*m feature arrays it was between 50-60 minutes and for 300 *μ*m feature arrays, about 90 minutes at 65°*C*. After the etching, the samples were rinsed in deionized water in order to stop the reaction, then dried under clean nitrogen and observed by optical and scanning electron microscopy (SEM). The optical microscopy gives an immediate evaluation of the etching homogeneity and microscopic defects present on the sample surface. Moreover, the feature base size, the undercut depth and the presence of any remaining SiO_2_ mask can also be estimated from optical microscopy. In order to measure the size and characterize the shape of pyramids, however, we needed to use SEM and to tilt the samples from normal incidence to angles from 40 to 70°. For SEM observation we used a tungsten-filament JEOL JSM-5910 microscope with a typical electron acceleration voltage of 20-30 kV.

After the observations, samples could be returned to the KOH solution in order to continue the etching to obtain the desired pin shape of the features. Several samples which were almost finished were placed into a ultrasonic bath in order to obtain sharp features by mechanical breaking of the remaining silicon at the top of the pin tip.

### Worm methods

The strains IG274^1^ and IG1061^13^ carrying the *frIS7* (*nlp-29p∷GFP; col-12p∷DsRed2*) integrated array in the wild-type or *sta-2(ok1860)* background, respectively were cultured under standard conditions, i.e. on *E. coli* OP50 on NGM agar plates^14^. Wounding with a micro-injection needle was performed as previously described^15^. For experiments using the silicon arrays, the wafers were mounted either on plastic handles using a cyanoacrylate adhesive (“superglue”) or on the end of syringe plungers, using household double-sided adhesive tape. Synchronized populations of young adult worms, obtained by alkaline hypochlorite treatment^14^, were transferred to plates without bacteria, rapidly wounded by bringing the wafer down with gentle pressure, repeatedly in a tiling fashion to cover the surface of the plate before transfer to a fresh OP50-seeded plate. In both cases, upregulation of *nlp-29p∷GFP* reporter gene levels were quantified after 6 hours with the COPAS Biosort (Union Biometrica)^15^. Staining of adult worms with Hoechst 33258 was performed as previously described^16^.

## Results

### Array microfabrication

In order to obtain an initial estimate of the required etching time and the final shape of the pins, we modelled the etching process using ACES software^17^, taking the data from Seidel *et al.*^18^. We simulated different mask shapes including circles (Fig. 3A) and squares (Fig. 3B) and we chose the former in order to simplify process optimization and to get closest to the desired final pin shape. The round features with initial mask diameters of 75, 100 and 300 *μm* were organized in arrays with separation distances sufficient for pyramid pin development. The pin density, i.e. the center-to-center pin distance, was chosen to be small enough to wound young adult *C. elegans* that are typically about 1 mm in length.

**Figure 3.**
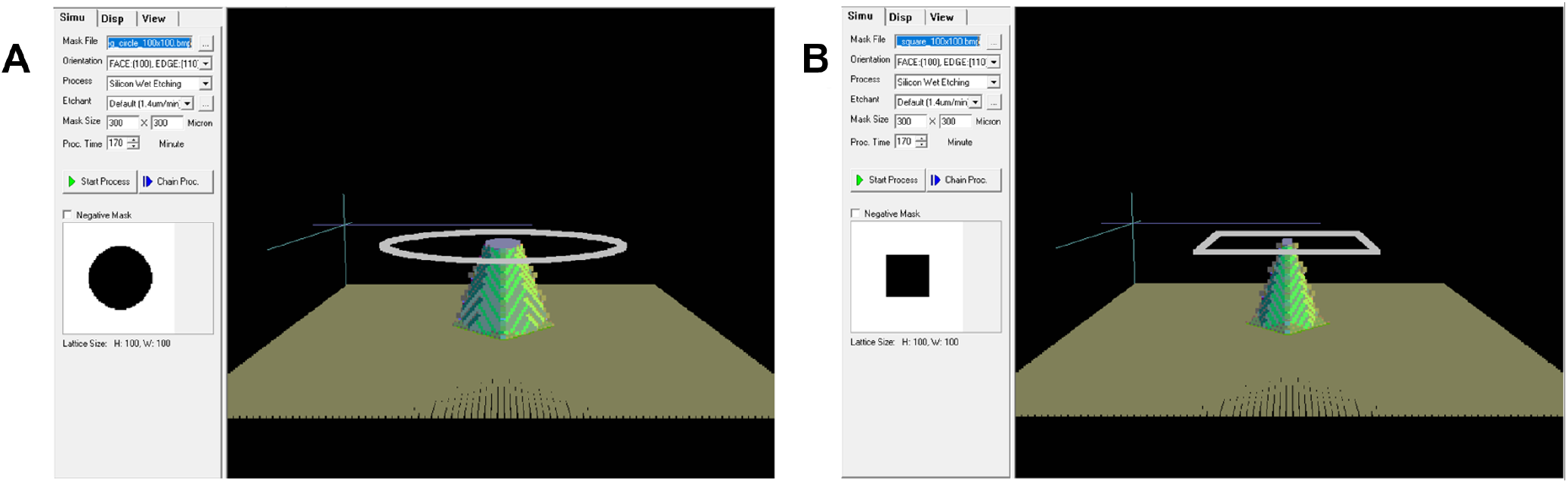
Simulation results for silicon (100) etching by KOH for round (A) and square (B) mask shapes.

The selectivity in the etching of Si and SiO_2_ was very high, and the oxide mask protected the silicon from alkaline attack relatively well. Fig. 4 shows the result of Si etching by KOH solution observed by SEM. KOH solution attacks (100) Si planes and, because of the strong undercut, the thin 290 nm-thick SiO_2_ layer remained only in an area with a size of 5 – 7*μm*. For some pyramids, we observed complete removal of the SiO_2_ protection, but in general, the etching was very homogeneous when we used our custom-designed 3D-printed rotating sample holder (Fig.2) at speeds of 100 to 180 rpm. SEM images similar to Fig. 4 were taken for all samples in order to ensure that etching was homogeneous over the entire wafer surface.

**Figure 4.**
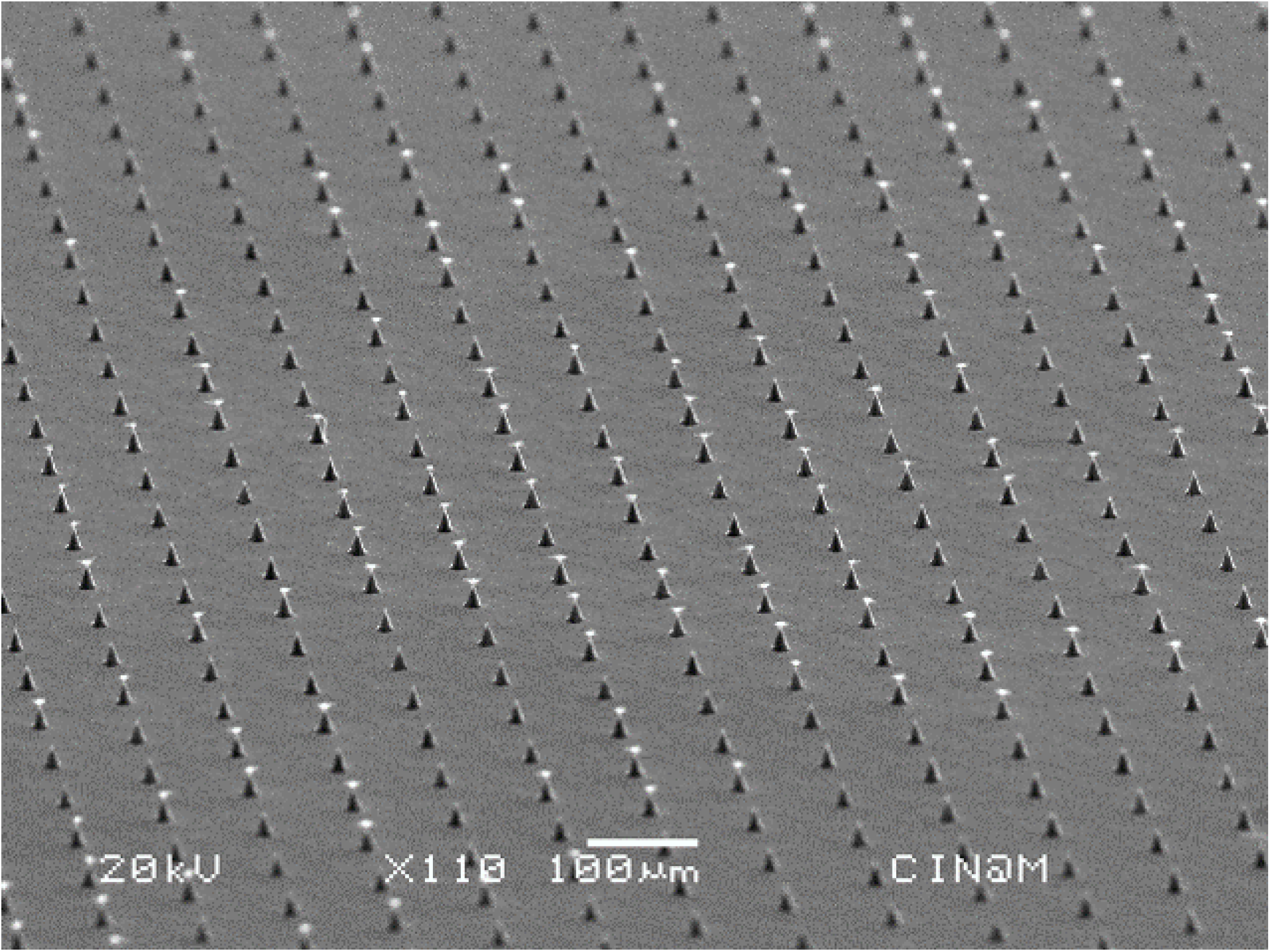
Large array of homogeneous pins at the end of the etching period. Of the 342 pins in this electron photomicrograph, 130 retain the “hat” that is subsequently removed in a pure water ultrasound bath. Manufacturing parameters: 75 *μ*m array; 750 ml fresh 45 % KOH; T=72C°; 100 rpm

Fig. 6 shows the successive SEM images of typical individual silicon pyramids that were taken on samples removed from KOH solution after 100, 110 and 118 minutes of etching. The solution was kept at the T=63°C and the rotation speed was 170 r.p.m. These pyramids were obtained using a hexagonal mask with the pitch of 100 *μ*m and initial feature diameter of 75 *μ*m. In the Fig. 6A obtained after 100 minutes of etching, one can clearly see the pyramid with a octagonal trunk, resulting from undercutting of silicon under the SiO_2_ mask, with a lateral size of about 30 *μ*m, corresponding to the pyramid base, and 16 *μ*m corresponding to the pyramid top. The remaining SiO_2_ mask is clearly visible on the top of the pyramid. Fig. 6B shows a pyramid obtained after 110 minutes of alkaline etching. One can see that the under-etching is deeper and the lateral sizes of both base and top of the pyramid are decreased to 25 and 5.5 *μ*m, respectively. Finally, Fig. 6C shows a pyramid obtained after etching in the same solution for a total time of 118 minutes. The remaining pyramid has a base size of 21 *μ*m and a very sharp tip. This form most closely resembles the pins previously described^2^.

In earlier work, the crystalline planes that form the pyramid pins were identified as (411)^3, 6, 19^. These crystal planes give a tetragonal tip similar to those observed for Si etching in Triton X-100 supplemented TMAH20. In our case, the tips of the pins are octagonal with the shapes similar to those reported by Wilke *et al.*^9, 21^. Those authors identified the crystal planes forming the main sharp part of pyramids as belonging to (312) family, while the crystalline planes of the base were identified as (228)^21^. In order to ensure scalability of the process, we compared the shape of the pyramids obtained using 75 and 100 *μ*m masks and found that the angles of the tips were similar; only the pin base size and height changed (Fig. 5). On the other hand, the shape of the tips was dependent on the etching bath conditions, in particular on the freshness of the KOH solution. Aging of the solution favored the formation of flat pyramid bases, consistent with previous observations^21^. Addition of ethanol, Triton X-100 or IPA also favored the formation of pyramids without sharp tips, formed by only their bases. Figure 7 shows successive SEM images of the pyramids etched at a temperature of 63°C in 45% KOH solution saturated with IPA. The images in Fig. 7 a-d correspond to etching times of 25, 50, 75 and 100 minutes, respectively. The addition of IPA stabilizes the etching process and pyramid surfaces are very smooth without defects. The crystal planes forming these pyramids correspond, however, to the pyramid bases and they are the same as observed in^21^.

**Figure 5.**
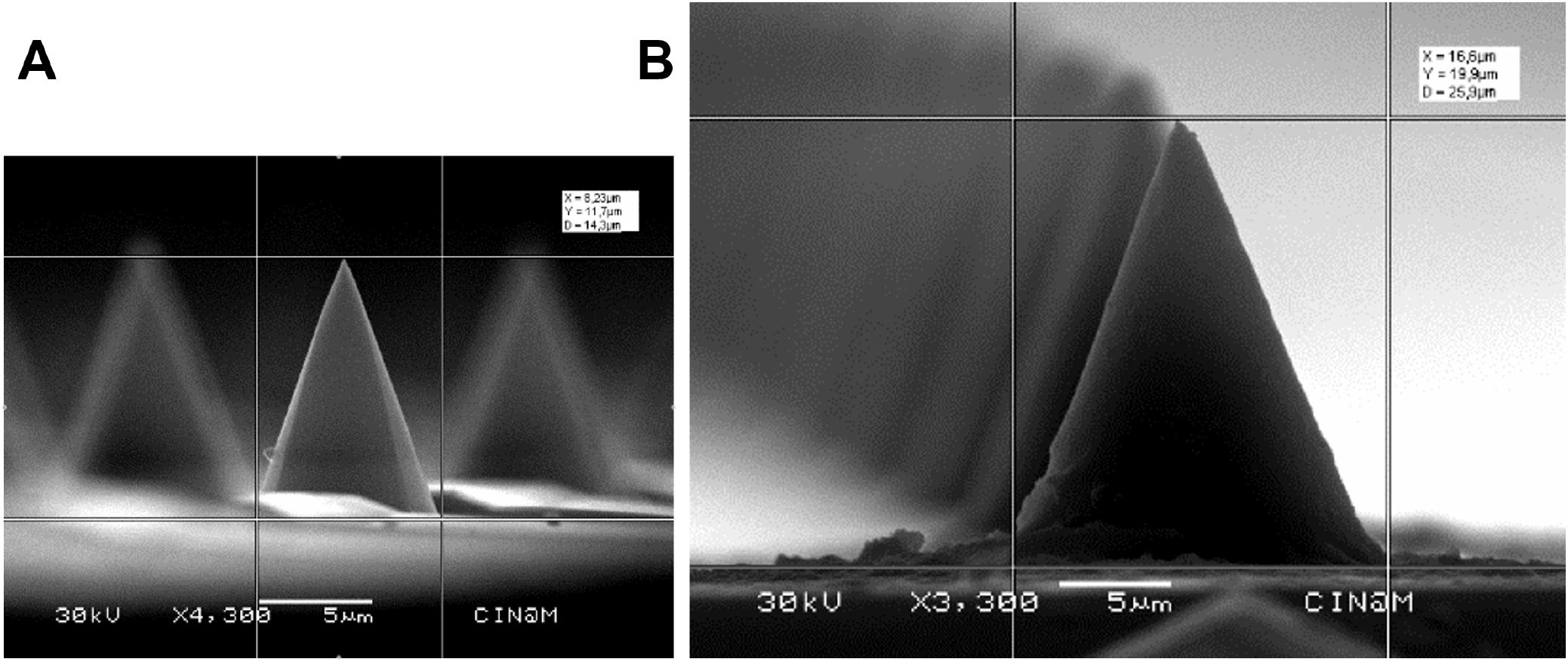
Silicon pyramids of different sizes retain a similar geometry. The small (A) and large (B) pyramids are of a similar shape, with the angles at the base being 110° and 115°, respectively. Manufacturing parameters: 75 *μ*m array; 39 min. (A); 100 *μ*m array; 70 min. (B); 400 ml fresh 45 % KOH; T=70C°; 160 rpm

**Figure 6.**
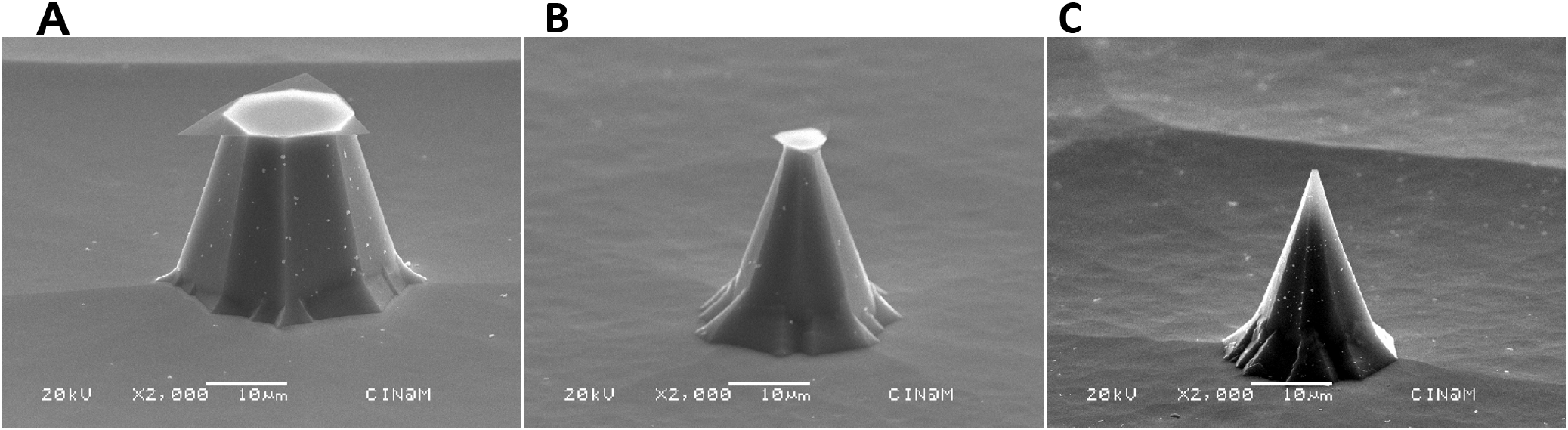
Silicon pyramids in 75 *μ*m array etched in KOH solution at T=63°C, sample holder rotation speed 170 r.p.m. Etching time: A) 100 minutes, B) 110 minutes C) 114 minutes (followed by an additional 4 minutes, after removal and inspection with a stereomicroscope). All the samples were tilted in the SEM at 70°.

**Figure 7.**
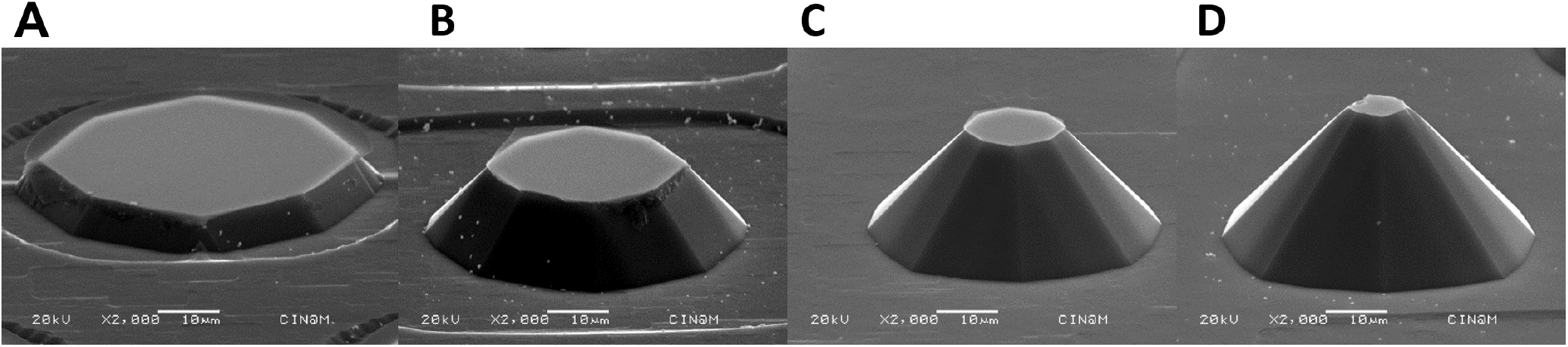
SEM images of silicon pyramids in 75 *μ*m array etched in IPA-saturated 45% KOH solution (44ml of IPA added to 400 ml KOH solution) at T=63° : A) 25 minutes, B) 50 minutes, C) 75 minutes, D) 100 minutes with agitation at 170 rpm. All the samples were tilted in the SEM 70°.

On the basis of these results, for subsequent etchings, we used only fresh aqueous KOH solution baths for each series of samples to obtain the optimal shape of the pyramidal pins and to diminish the base development. We also increased the etching temperature because higher temperatures allowed us to decrease the etching time which was otherwise inconveniently long for the arrays with initial sizes of 100 and 300 *μ*m. For 75 *μ*m arrays the typical etching time was about 45 minutes for standard etching temperatures of 70-72°. In Fig. 8A we show a pyramid obtained after 44 minutes of KOH etching at 70°C. One can clearly see the pyramid covered by the remaining SiO_2_ protection layer, with a more extensive fine, electron-transparent SiO_2_ “veil”. The octagonal shape of the pyramid tip is also very clearly visible. Fig. 8B) shows a pyramid obtained after 45 minutes of etching. The pyramid tip is much sharper than the previous one. The fine electron-transparent oxide layer lost its mechanical stability and was removed during the rinsing and drying of the sample. In order to sharpen the tips we returned the same sample to the KOH etching solution for 3 supplementary minutes (Fig. 8C). The sharp pyramids were rapidly attacked by KOH and when over-etched, only the more resistant crystalline planes close to the base remained (Fig. 8C). The height as well as width of pyramids diminished very rapidly in KOH solution when the SiO_2_ protection was gone. We obtained similar results for a sample etched for 47 minutes in one run without removal from the solution (Fig. 8D). This instability made the etching process very time-sensitive. We therefore etched samples in KOH for a time slightly inferior to that expected for complete SiO_2_ removal, then examined the features using SEM. On the basis of the extent of etching, if necessary we then returned the sample to the KOH solution to finish the process. Because of the consumption of all the silicon under the SiO_2_ mask, the “hat” part is completely removed and washed out by KOH solution or while the wafer is rinsed under a flow of deionized water after the etching process has finished. When the pyramid tip size approached 2-5 *μ*m, it was also possible to finish the fabrication by mechanical breaking of the remaining “hat” in a pure water ultrasound bath. In both cases, the remaining pyramids have very sharp tips.

**Figure 8.**
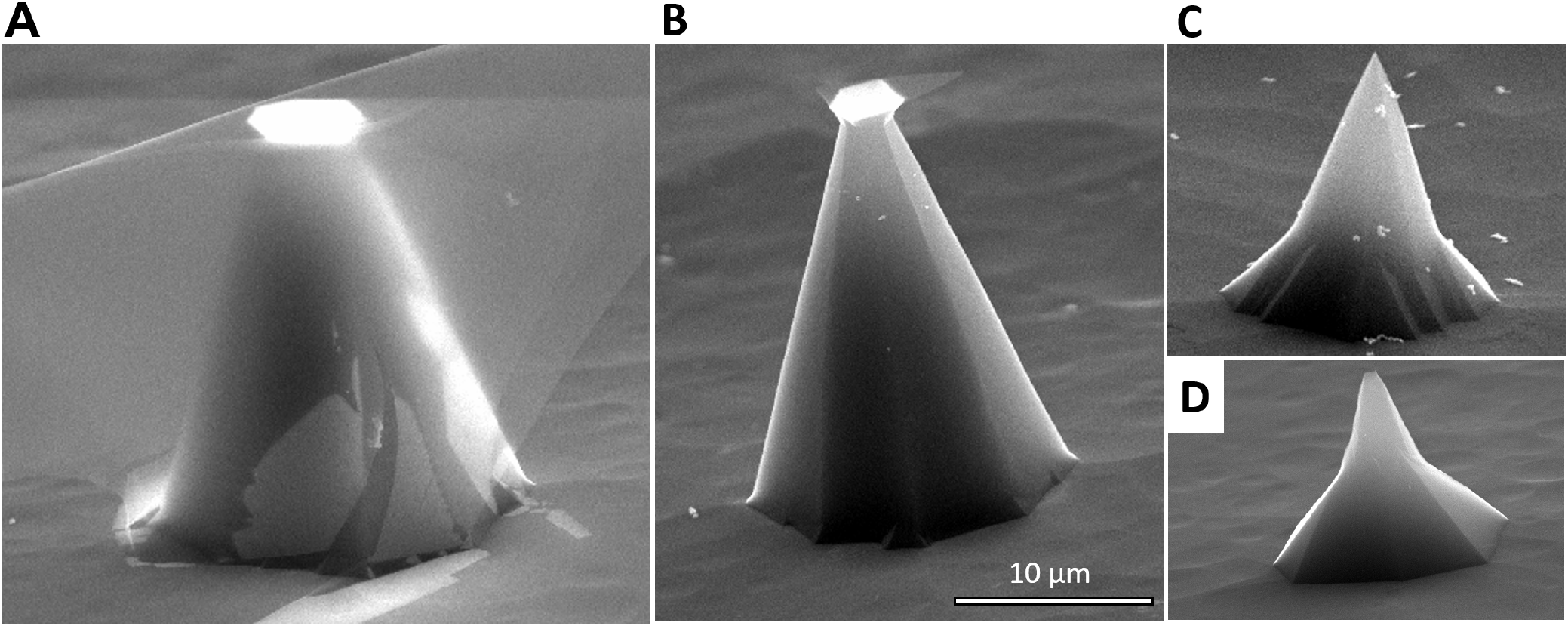
SEM images of silicon pyramid needles etched in 45% KOH solution at T=72°, sample holder rotation speed 170 r.p.m.: A) 44 minutes B) 45 minutes C) 45 (followed by an additional 3 minutes, after removal and inspection with a stereomicroscope) D) 47 minutes (continuous). All the samples were tilted in the SEM 70°.

### Efficient wounding of *C. elegans*

One of the consequences of wounding or fungal infection of the *C. elegans* epidermis is the rapid induction of antimicrobial peptide (AMP) gene expression. This can be monitored through the use of transgenic worms carrying fluorescent reporter genes. The most extensively characterised strain, IG274, carries the integrated transgene *frIs7*, with 2 reporter genes, dsRed and GFP under the control of a constitutive epidermal promoter, *col-12*, and the infection-inducible *nlp-29* promoter, respectively. In the absence of pathogens or injury, IG274 worms appear red, but express GFP strongly when wounded or infected. These changes can be monitored visually or quantified using the COPAS Biosort^1^. In a first series of experiments, we compared the induction of *nlp-29p∷GFP* provoked by wounding by hand with a glass needle or with the silicon pin arrays. Arrays were adhered to a short plastic rod for ease of handling and brought down gently onto a population of worms on an agar plate. The array was lifted, moved and brought down again. The light pattern of indentations left on the surface of the agar was used to ensure complete coverage of the population. As expected, manual wounding with a microinjection needle caused a substantial increase in average GFP expression, and this was almost completely abrogated in a *sta-2* mutant, consistent with previous observations^13^. There was an even more marked increase when worms, whether at a lower or higher density on the culture plate, were wounded with a silicon array. Again, this response was dependent upon *sta-2*, consistent with both methods activating AMP gene expression by the same canonical signal transduction pathway (Figure 9A). Importantly, in addition to producing equivalent results as the standard needle method, the use of the array simplified and accelerated the procedure substantially. An experienced researcher with good manual dexterity can reasonably wound one worm every 2-4 seconds. With the arrays, no training was required and the speed of wounding was increased at least 10 fold, as hundreds of worms could be wounded in less than 30 seconds. We found that we could use the arrays multiple times. After repeated use, in some cases, we observed a slight abrasion of the pin apex, presumably because of strong mechanical pressure of the tips against the agar support (Fig. 10). To demonstrate that the arrays truly pierced the worms, we made use of the fact that *C. elegans* is surrounded by a collagen-rich cuticle that renders it impermeable to most compounds^22^. Under normal culture conditions, in non-molting animals, the cell membrane permeable Hoechst 33258 dye can only stain the nuclei of intestinal cells. Mutants with a compromised epithelial barrier on the other hand display staining of epidermal nuclei^23^. In clear contrast to control worms, we observed Hoechst staining of epidermal nuclei in the population of wounded worms (Figure 9B). Having established the potential utility of the arrays, we went on to evaluate the efficiency of wounding using pins of different sizes, spacing and geometries. While the average fluorescence ratio was greater when the pins were longer, the geometry did not appear to be critical (Figure 11).

**Figure 9.**
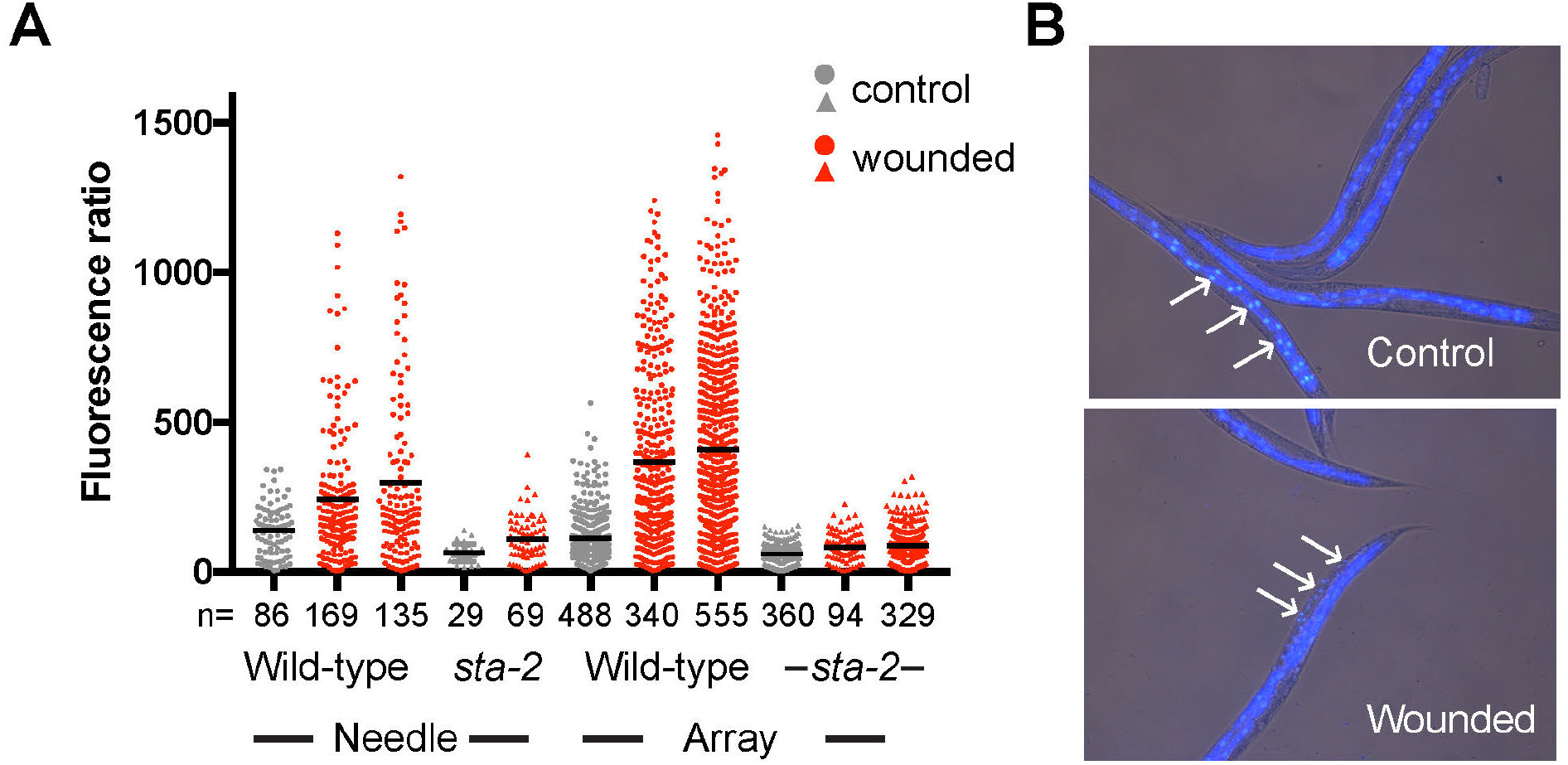
Efficient wounding of *C. elegans* with a silicon array. **A.** Quantification of the change of reporter gene expression provoked by wounding with a needle (5 left-hand columns), or arrays, worms carrying *frIs7* in the wild-type or *sta-2* mutant background (circles and triangles, respectively). The fluorescence ratio (Green/TOF in arbitrary but constant units) is shown for the indicated number of worms for each condition (n). The bars indicate the means. **B.** Representative images of control (top) and wounded (bottom) adult wild-type worms stained with Hoechst 33258. In the control animals, only staining of the intestinal nuclei is apparent (arrows), while in the array-wounded worms nuclei in the epidermis (arrows) are also stained.

**Figure 10.**
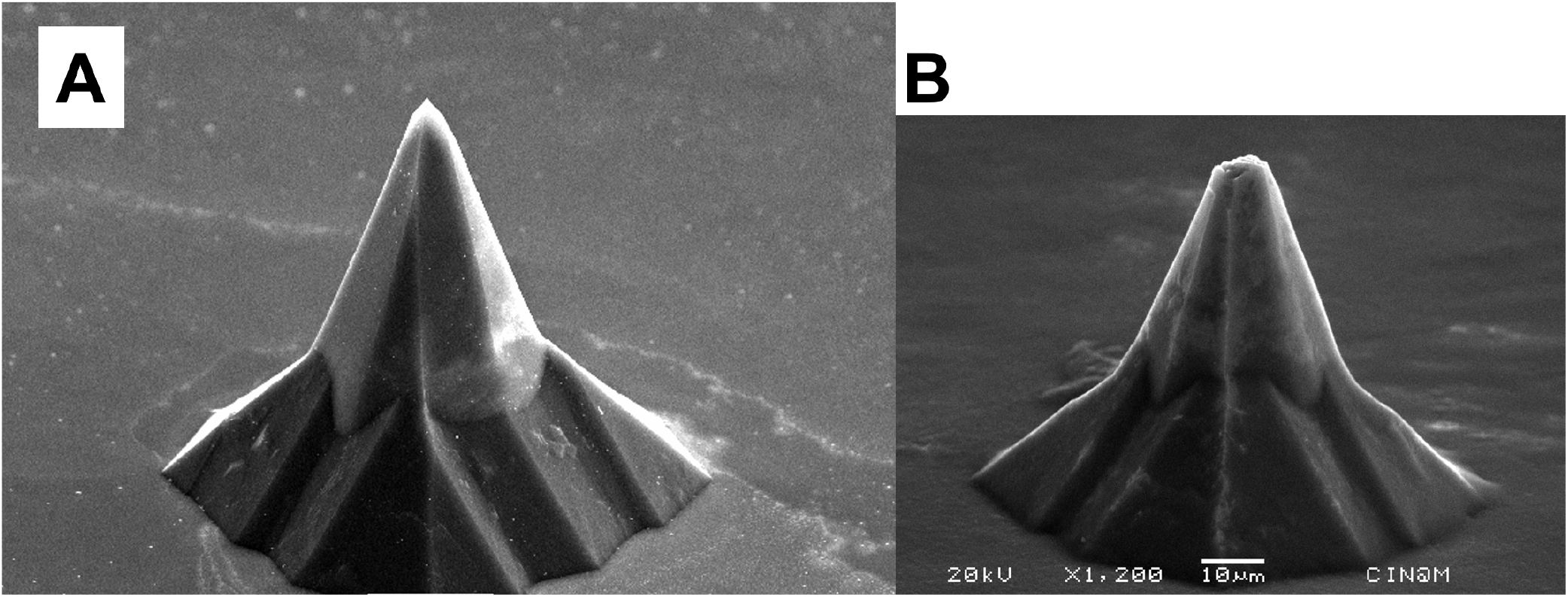
SEM images of silicon pyramid needles A) before and B) after worm experiments

**Figure 11.**
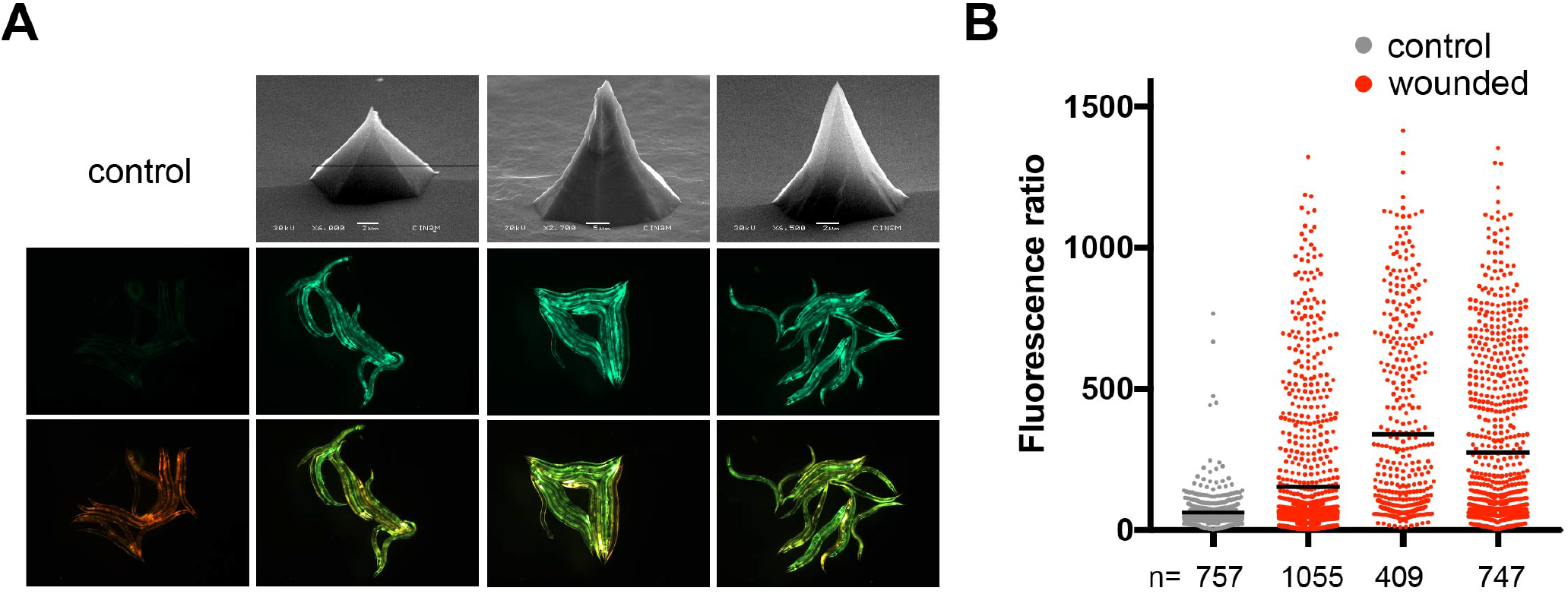
Wounding of *C. elegans* with different silicon arrays. **A.** Top row shows SEM images of 3 different arrays (left to right: H75-42, H100-9, H75-46). Note that, as indicated, the magnification is not the same for all 3, hence the scale bars are different too. The other 2 rows are GFP (middle row) or mixed GFP/dsRed fluorescence signals for representative groups of worms carrying *frIs7* picked because they appeared to have a normal morphology (i.e. were not badly wounded) after treatment with the indicated array, compared to non-wounded controls (left-hand column). **B.** Quantification of the change of reporter gene expression provoked by wounding with the 3 arrays, in the same order as in A, in worms carrying *frIs7*. Details are as in the previous figure.

## Discussion

Using an alkaline etching technique, we produced arrays of silicon pyramidal pins. Contrary to the fabrication of concave features on Si (100) which is usually self-limiting when the slowly etched (111) surfaces are reached, the fabrication of convex features on Si (100) by alkaline etching is more challenging^5^. Two processes occur at the same time: etching of the (100) crystalline surface directed towards the wafer depth in the areas that are not protected by a mask and undercutting of silicon features in a lateral direction under the mask despite the protection. The final shape of the obtained features depends on the difference in the rates of these two processes^24^. For example, the etching rate for (111) silicon crystalline planes is more than 100 times slower than for the other planes such as (100)^25, 26^, and it is possible to obtain sharp features with a pyramidal shape both for (100) and (111) oriented wafers^20^. Numerous models have been proposed to explain the convex corner undercutting^25^ and its compensation^26^ and, more generally, anisotropic etching of silicon^27^. The main complexity in the description of experimental procedures arises from the fact that multiple parameters have been reported to affect the undercutting rate: the solution temperature^5^, KOH or TMAH concentration, freshness of the alkaline solution^21^ as well as the introduction of different chemical additives to the solutions such as I_2_/KI^28^, the surfactant Triton-X-100^20, 29^ or IPA^5^. In our fabrication process, we used only fresh aqueous KOH solution baths for each sample series and we optimized the shape of the pyramidal pins to be suitable for biological experiments. Only etching in pure KOH solutions allowed us to obtain sharp pyramids. Addition of ethyl or isopropyl alcohol or Triton to aqueous KOH solutions favoured the formation of crystal planes forming flat pyramids. We developed a series of sample holders for the wafers of different sizes from 1cm^2^ up to 100 mm (4 inches) resistant to KOH and alcohol solutions up to the temperatures of more than 80°C necessary to ensure homogeneous etching over large surfaces. Very sharp pyramidal needles were organized into arrays with a pitch of 100 - 133 *μ*m, dense enough to wound young adult *C. elegans* hermaphrodites that are around 1 mm in length. In the original publication, piercing structures similar to the ones described here were claimed to have been used to transform the entomopathogenic nematode *Heterorhabditis bacteriophora* in an extremely rapid and efficient manner^2^. Despite the obvious attraction for such a simple method, no other publication has ever reported the successful replication of this work. Parasitic nematodes are notoriously difficult to transform, compared to *C. elegans* for which micro-injection-based transformation is a standard laboratory technique. In our hands, despite numerous attempts, we were unable to obtain any evidence for transformation of *C. elegans* using our silicon pin arrays (JB and CC, unpublished). Thus, unfortunately, it appears that their utility for making transgenic nematodes is limited at best. On the other hand, these arrays enormously accelerate and simplify the generation of large populations of wounded *C. elegans*. This opens up the potential for generating samples in a sufficient volume to allow transciptomic, proteomic or metabolomic analyses of the response to wounding, further strengthening the utility of *C. elegans* in this domain^30^.

## Acknowledgements

Supported by institutional grants from the Institut national de la santé et de la recherche médicale, Centre national de la recherche scientifique and Aix-Marseille University to the CIML, and the Agence Nationale de la Recherche program grants (ANR-12-BSV3-0001-01, ANR-16-CE15-0001-01, ANR-11-LABX-0054 (Labex INFORM) and ANR-11-IDEX-0001-02 (A*MIDEX)) to JJE. This work was partly supported by the French RENATECH network. Microfabrication and electron microscopy were performed in PLANETE CT PACA clean room facility. Worm sorting was performed using the facilities of the French National Functional Genomics platform, supported by the GIS IBiSA and Labex INFORM. The authors thank Shizue Omi for worm optical imaging, as well as Aicha Aouane and Fabrice Richard from IBDM for preparation of worms for electron microscopy.

## Author contributions statement

J.B., I.O and J.J.E. conceived the experiments; J.B., C.C., I.O and F.B. conducted pin array microfabrication; F.B. and I.O. the SEM observations; J.B., C.C. and J.J.E. the worm experiments; I.O. and J.J.E wrote the manuscript; All authors contributed to the discussion and reviewed the manuscript.

